# An orally bioavailable broad-spectrum antiviral inhibits SARS-CoV-2 and multiple endemic, epidemic and bat coronavirus

**DOI:** 10.1101/2020.03.19.997890

**Authors:** Timothy P. Sheahan, Amy C. Sims, Shuntai Zhou, Rachel L. Graham, Collin S. Hill, Sarah R. Leist, Alexandra Schäfer, Kenneth H. Dinnon, Stephanie A. Montgomery, Maria L. Agostini, Andrea J. Pruijssers, James D. Chapell, Ariane J. Brown, Gregory R. Bluemling, Michael G. Natchus, Manohar Saindane, Alexander A. Kolykhalov, George Painter, Jennifer Harcourt, Azaibi Tamin, Natalie J. Thornburg, Ronald Swanstrom, Mark R. Denison, Ralph S. Baric

**Affiliations:** Department of Epidemiology, University of North Carolina at Chapel Hill, Chapel Hill, NC; Lineberger Comprehensive Cancer Center, University of North Carolina at Chapel Hill, Chapel Hill, NC; Department of Pathology & Laboratory Medicine, University of North Carolina, Chapel Hill, NC; Department of Pediatrics, Vanderbilt University Medical Center, Nashville, TN; Emory Institute of Drug Development (EIDD), Emory University, Atlanta, GA; Drug Innovation Ventures at Emory (DRIVE), Atlanta, GA; Department of Pharmacology and Chemical Biology, Emory University, Atlanta, GA; Centers for Disease Control and Prevention, Division of Viral Diseases Atlanta GA; Department of Biochemistry and Biophysics, University of North Carolina at Chapel Hill, Chapel Hill, NC; Department of Microbiology and Immunology, University of North Carolina at Chapel Hill, Chapel Hill, NC

## Abstract

Coronaviruses (CoVs) traffic frequently between species resulting in novel disease outbreaks, most recently exemplified by the newly emerged SARS-CoV-2. Herein, we show that the ribonucleoside analog β-D-N^4^-hydroxycytidine (NHC, EIDD-1931) has broad spectrum antiviral activity against SARS-CoV 2, MERS-CoV, SARS-CoV, and related zoonotic group 2b or 2c Bat-CoVs, as well as increased potency against a coronavirus bearing resistance mutations to another nucleoside analog inhibitor. In mice infected with SARS-CoV or MERS-CoV, both prophylactic and therapeutic administration of EIDD-2801, an orally bioavailable NHC-prodrug (b-D-N^4^-hydroxycytidine-5’-isopropyl ester), improved pulmonary function, and reduced virus titer and body weight loss. Decreased MERS-CoV yields *in vitro* and *in vivo* were associated with increased transition mutation frequency in viral but not host cell RNA, supporting a mechanism of lethal mutagenesis. The potency of NHC/EIDD-2801 against multiple coronaviruses, its therapeutic efficacy, and oral bioavailability *in vivo*, all highlight its potential utility as an effective antiviral against SARS-CoV-2 and other future zoonotic coronaviruses.

## Introduction

The genetically diverse *Orthocoronavirinae* (coronavirus, CoV) family circulates in many avian and mammalian species. Phylogenetically, CoVs are divided into 4 genera: alpha (group 1), beta (group 2), gamma (group 3) and delta (group 4). Three new human CoV have emerged in the past 20 years with severe acute respiratory syndrome CoV (SARS-CoV) in 2002, Middle East respiratory syndrome CoV (MERS-CoV) in 2012, and now SARS-CoV-2 in 2019^1–3^. The ongoing SARS-CoV-2 epidemic (referred to as COVID-19, Coronavirus disease 2019) has caused over 89,000 infections and over 3,000 deaths in 71 countries. Like SARS- and MERS-CoV, the respiratory disease caused by SARS-CoV-2 can progress to acute lung injury (ALI), an end stage lung disease with limited treatment options and very poor prognoses^3–5^. This emergence paradigm is not limited to humans. A novel group 1 CoV called swine acute diarrhea syndrome CoV (SADS-CoV) recently emerged from bats causing the loss of over 20,000 pigs in Guangdong Province, China^6^. More alarmingly, many group 2 SARS-like and MERS-like coronaviruses are circulating in bat reservoir species that can use human receptors and replicate efficiently in primary human lung cells without adaptation^6–9^. The presence of these “pre-epidemic” zoonotic strains foreshadow the emergence and epidemic potential of additional SARS-like and MERS-like viruses in the future. Given the diversity of CoV strains in zoonotic reservoirs and a penchant for emergence, broadly active antivirals are clearly needed for rapid response to new CoV outbreaks in humans and domesticated animals.

Currently, there are no approved therapies specific for any human CoV. β-D-N4-hydroxycytidine (NHC, EIDD-1931) is orally bioavailable ribonucleoside analog with broad-spectrum antiviral activity against various unrelated RNA viruses including influenza, Ebola, CoV and Venezuelan equine encephalitis virus (VEEV)^10–13^. For VEEV, the mechanism of action (MOA) for NHC has been shown to be through lethal mutagenesis where deleterious transition mutations accumulate in viral RNA^11,14^. Here, we demonstrate that NHC exerts potent, broad-spectrum activity against SARS-CoV, MERS-CoV and their related bat-CoV in primary human airway epithelial cell cultures (HAE), a biologically relevant model of the human conducting airway. In addition, we show that NHC is potently antiviral against the newly emerging SARS-CoV-2 as well as against coronavirus bearing resistance mutations to the potent nucleoside analog inhibitor, remdesivir (RDV). In SARS- or MERS-CoV infected mice, both prophylactic and therapeutic administration EIDD-2801, an oral NHC-prodrug (b-D-N^4^-hydroxycytidine-5’-isopropyl ester) improved pulmonary function and reduced virus titer and ameliorated disease severity. In addition, therapeutic EIDD-2801 reduced the pathological features of ALI in SARS-CoV infected mice. Using a high-fidelity deep sequencing approach (Primer ID), we found that increased mutation rates coincide with decreased MERS-CoV yields *in vitro* and protective efficacy *in vivo* supporting the MOA of lethal mutagenesis against emerging CoV^13^. The broad activity and therapeutic efficacy of NHC/EIDD-2801 highlight its potential to diminish epidemic disease today and limit future emerging CoV outbreaks.

## Results

### NHC potently Inhibits MERS-CoV, SARS-CoV and newly emerging SARS-CoV-2 Replication

To determine whether NHC blocks the replication of highly pathogenic human CoV, we performed antiviral assays in continuous and primary human lung cell cultures. We first assessed the antiviral activity of NHC against MERS-CoV in the human lung epithelial cell line Calu-3 2B4 (“Calu3” cells). Using a recombinant MERS-CoV expressing nanoluciferase (MERS-nLUC)^15^, we measured virus replication in cultures exposed to a dose range of drug for 48hr. NHC was potently antiviral with an average half-maximum effective concentration (IC_50_) of 0.15μM and no observed cytoxicity in similarly treated uninfected cultures across the dose range (50% cytotoxic concentration, CC50, >10μM) (Fig. 1A). The therapeutic index for NHC was >100. Similarly, NHC strongly inhibited SARS-CoV-2 replication in Vero cells with an IC50 of 0.3μM and CC50 of >10μM (Fig. 1B). Human primary airway epithelial (HAE) cell cultures model the architecture and cellular complexity of the conducting airway and are readily infected by multiple human and zoonotic CoV, including SARS- and MERS-CoV^16^. We first assessed cytotoxicity of NHC in HAE treated with an extended dose range for 48hr using quantitative PCR of cell death-related gene transcripts as our metric. NHC treatment did not appreciably alter gene expression even at doses up to 100μM (Supplementary Figure 1). In MERS-CoV infected HAE, NHC dramatically reduced virus production with maximal titer reduction of > 5 logs at 10μM (average IC_50_ = 0.024 μM), which correlated with reduced genomic (ORF1) and subgenomic (ORFN) RNA in paired samples (Fig. 1C). We observed similar trends in titer reduction (> 3 log at 10μM, average IC_50_ = 0.14 μM) and levels of genomic and subgenomic RNA in SARS-CoV infected HAE (Fig. 1D). Thus, NHC was potently antiviral against MERS-CoV and SARS-CoV-2 in cell lines and MERS-CoV and SARS-CoV in human primary HAE cell cultures without cytotoxicity.

**Figure 1:**
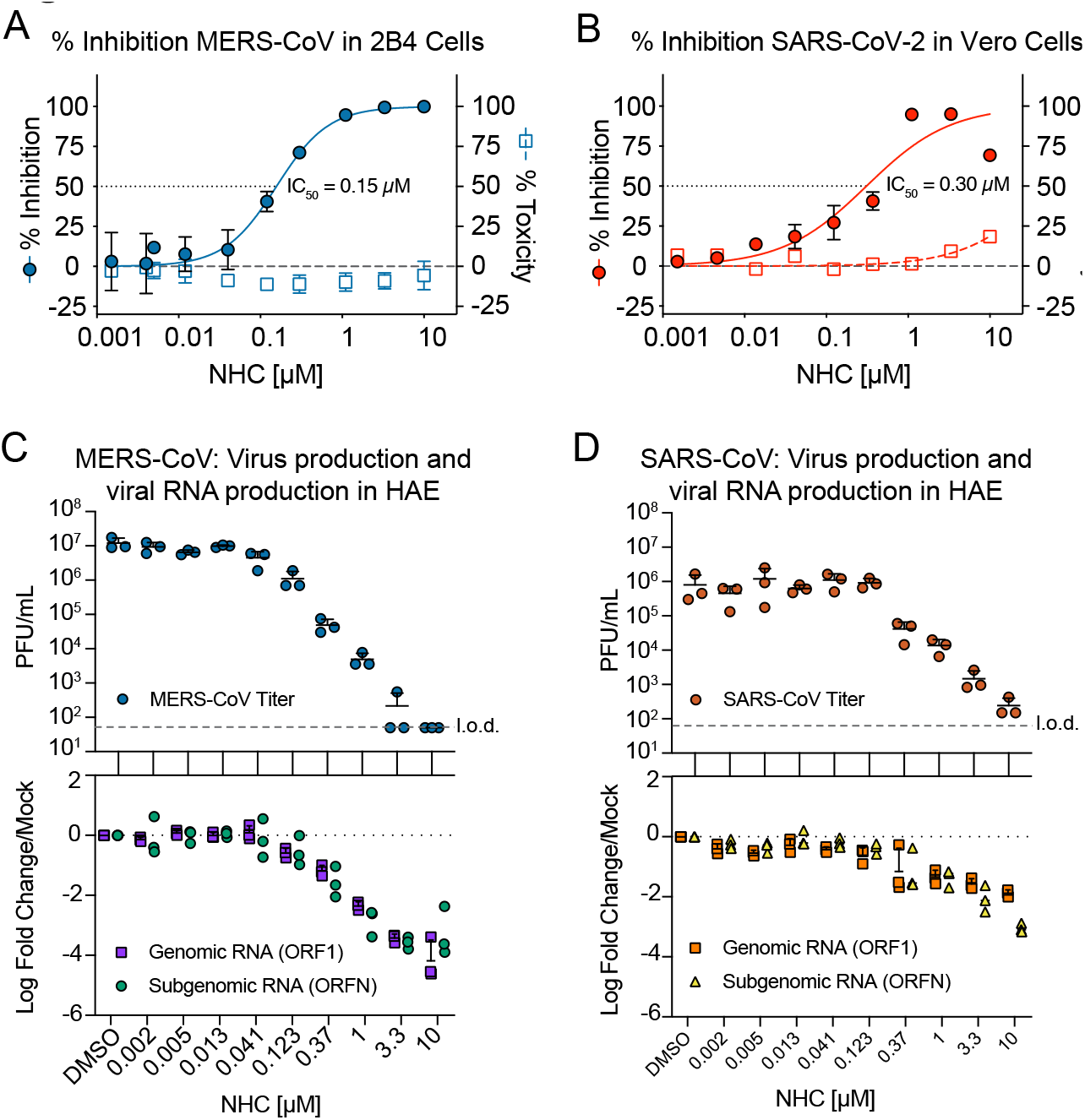
NHC potently Inhibits MERS-CoV, SARS-CoV and newly emerging SARS-CoV-2 Replication. **a**, NHC antiviral activity and cytotoxicity in Calu3 cells infected with MERS-CoV. Calu3 cells were infected in triplicate with MERS-CoV nanoluciferase (nLUC) at a multiplicity of infection (MOI) of 0.08 in the presence of a dose response of drug for 48 hours, after which replication was measured through quantitation of MERS-CoV–expressed nLUC. Cytotoxicity was measured in similarly treated but uninfected cultures via Cell-Titer-Glo assay. Data is combined from 3 independent experiments. **b**, NHC antiviral activity and cytotoxicity in Vero cells infected with SARS-CoV-2. Vero cells were infected in duplicate with SARS-CoV-2 clinical isolate virus at an MOI of 0.05 in the presence of a dose response of drug for 48 hours, after which replication was measured through quantitation of cell viability by Cell-Titer-Glo assay. Cytotoxicity was measured as in **a**. Data is combined from 2 independent experiments. **c**, NHC inhibits MERS-CoV virus production and RNA synthesis in primary human lung epithelial cell cultures (HAE). HAE cells were infected with MERS-CoV red fluorescent protein (RFP) at an MOI of 0.5 in duplicate in the presence of NHC for 48 hours, after which apical washes were collected for virus titration. qRT-PCR for MERS-CoV ORF1 and ORFN mRNA. Total RNA was isolated from cultures in **c** for qRT-PCR analysis. Representative data from three separate experiments with three different cell donors are displayed. PFU, plaque-forming units. **d**, NHC inhibits SARS-CoV virus production and RNA synthesis in primary human lung epithelial cell cultures (HAE). Studies performed as in **c** but with SARS-CoV green fluorescent protein (GFP). Representative data from two separate experiments with two different cell donors are displayed.

### NHC is effective against remdesivir resistant virus and multiple distinct zoonotic CoV

All human CoV are thought to have emerged as zoonoses most recently exemplified by SARS-CoV, MERS-CoV and SARS-CoV-2^17–19^. Although taxonomically divided into multiple genogroups (alpha, beta, gamma, delta), human CoV are found in only the alpha and beta subgroups thus far (Fig. 2A). There is high sequence conservation in the RdRp across CoV (Fig. 2A). For example, the RdRp of SARS-CoV-2 has 99.1% similarity and 96% amino acid identity to that of SARS-CoV (Fig. 2A). To gain insight into structural conservation of RdRp across the CoV family, we modeled the variation reflected in the RdRp dendrogram in Fig. 2A onto the structure of the SARS-CoV RdRp^20^ (Fig. 2B). The core of the RdRp molecule and main structural motifs that all RdRp harbor (Fig. 2B and Supplementary Figure 2) are highly conserved among CoV including SARS-CoV-2. We previously reported that CoV resistance to another broad spectrum nucleoside analog, remdesivir (RDV) was mediated by RdRp residues F480L and V557L in a model coronavirus mouse hepatitis virus (MHV) and in SARS-CoV, resulting in a 5-fold shift in IC_50_ (Fig. 2C)^21^. Consequently, we tested whether RDV resistance mutations in MHV conferred cross resistance to NHC (Figure 2D). In fact, the two RDV resistance mutations, alone or together conferred increased sensitivity to inhibition by NHC. As our previous studies have demonstrated a high genetic barrier to NHC for VEEV, influenza and coronavirus^11–13^, the lack of cross resistance further suggests that NHC and RDV may select for exclusive and mutually sensitizing resistance pathways.

**Figure 2.**
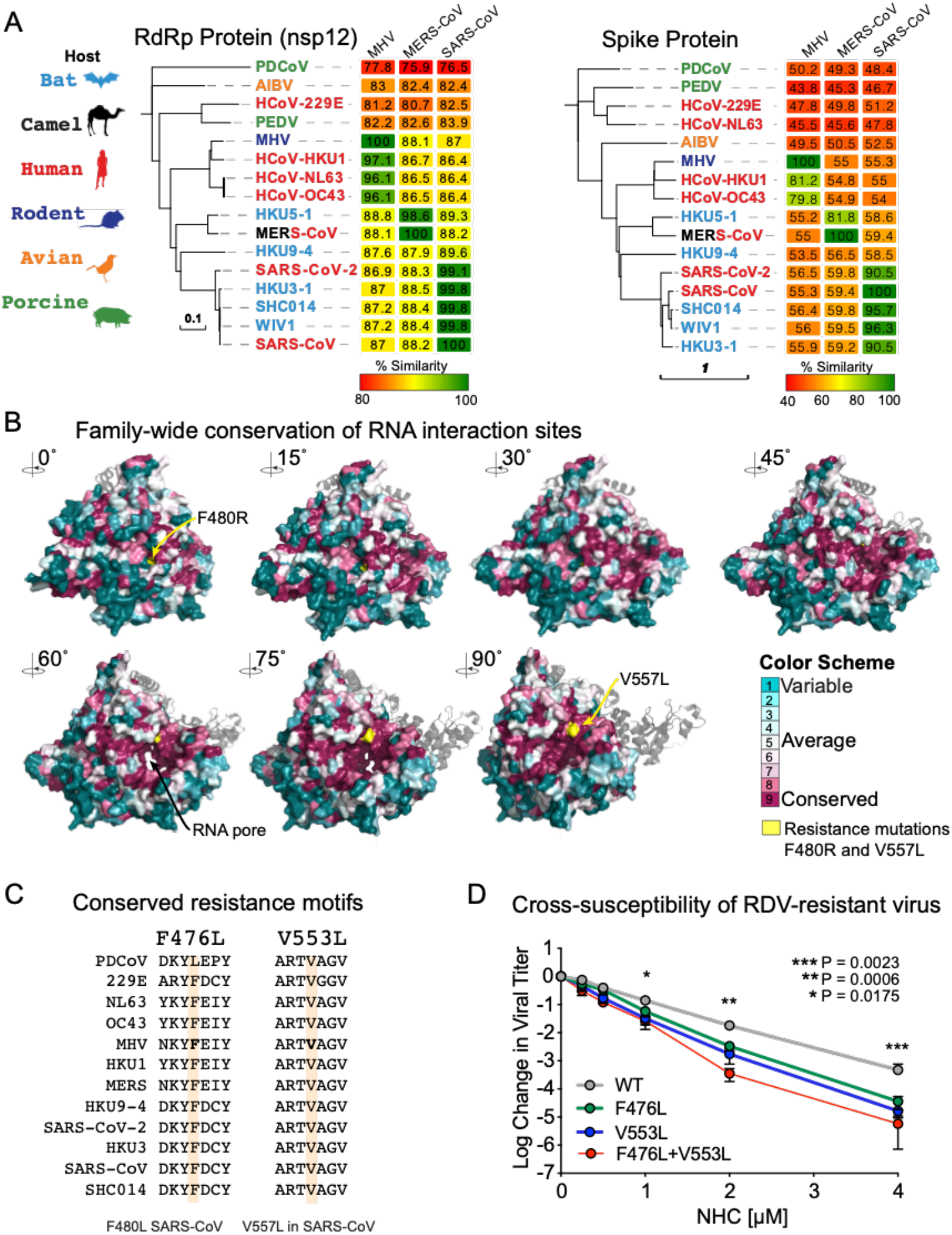
Remdesivir resistance mutations in the highly conserved RNA-dependent RNA polymerase increase susceptibility to NHC. **a**, Neighbor-joining trees created with representatives from all four CoV genogroups showing the genetic similarity of CoV nsp12 (RdRp) and CoV spike glycoprotein, which mediates host tropism and entry into cells. Text color of the virus strain label corresponds to virus host species on the left. The heatmap adjacent to each neighbor-joining tree depicts percent amino acid identity (% A.A. similarity) against mouse hepatitis virus (MHV), SARS-CoV or MERS-CoV. **b**, Core residues of the CoV RdRp are highly conserved among CoV. The variation encompassed in panel **a** was modeled onto the RdRp structure of the SARS-CoV RdRp. **c**, Amino acid sequence of CoV in panel **a** at known resistance alleles to antiviral drug remdesivir (RDV). **d**, RDV resistant viruses are more susceptible to NHC antiviral activity. Virus titer reduction assay across a dose response of NHC with recombinant MHV bearing resistance mutations to RDV. Asterisks indicate statistically significant differences by Mann-Whitney test.

To explore the breadth of antiviral efficacy against zoonotic CoV, we performed antiviral assays in HAE with three zoonotic Bat-CoV, SHC014, HKU3 and HKU5. Closely related to the beta 2b SARS-CoV, Bat-CoV SHC014 is capable of replicating in human cells without adaptation^8^ suggesting its potential for zoonotic emergence. More distantly related SARS-like beta 2b CoV, recombinant Bat-CoV HKU3 has a modified receptor binding domain to facilitate growth in cell culture^22^. Lastly, Bat-CoV HKU5 is a MERS-like beta 2c CoV^23^. NHC diminished infectious virus production and the levels of genomic/subgenomic viral RNA in HAE in a dose-dependent manner for all three Bat-CoVs (Fig. 3). Therefore, the antiviral activity of NHC was not limited by natural amino acid variation in the RdRp, which among the group 2b and group 2c CoV can vary by almost 20% (Fig. 2A). Moreover, these data suggest that if another SARS- or MERS-like virus were to spillover into humans in the future, they would likely be susceptible to the antiviral activity of NHC.

**Figure 3:**
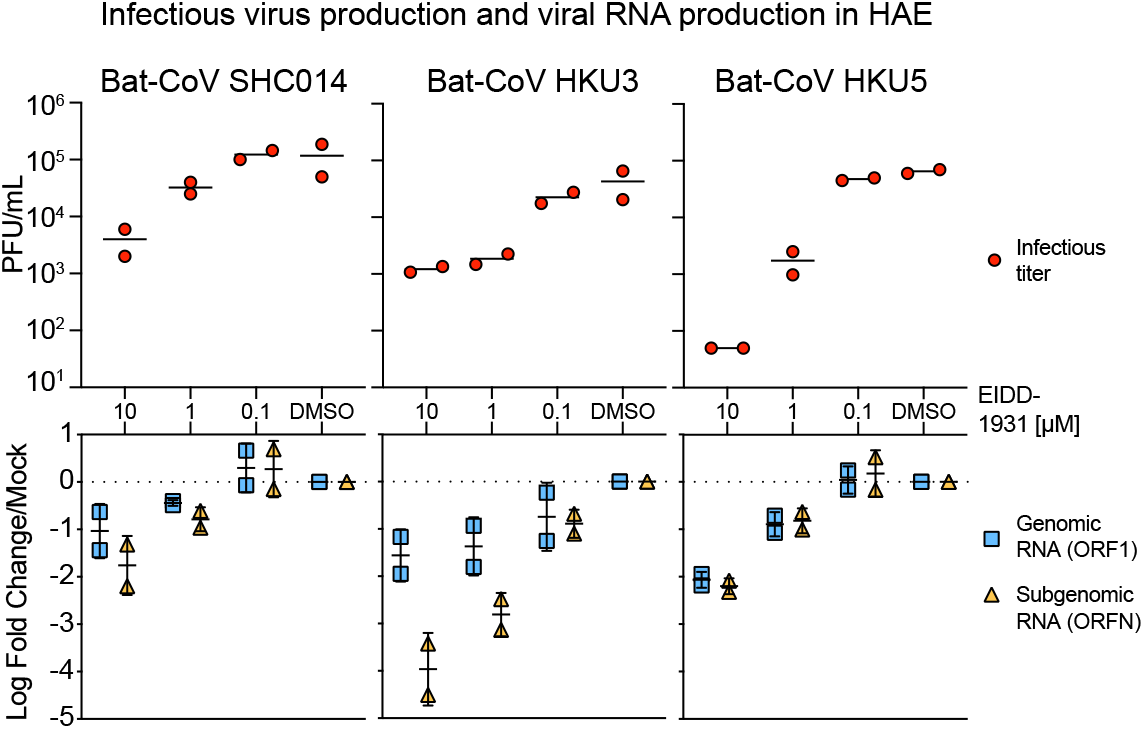
NHC is effective against multiple genetically distinct Bat-CoV. Top: Antiviral efficacy of NHC in HAE cells against SARS-like (HKU3, SHC014, group 2b) and MERS-like (HKU5, group 2c) bat-CoV. HAE cells were infected at an MOI of 0.5 in the presence of NHC in duplicate. After 48 hours, virus produced was titrated via plaque assay. Each data point represents the titer per culture. Bottom: qRT-PCR for CoV ORF1 and ORFN mRNA in total RNA from cultures in the top panel. Representative data from two separate experiments with two different cell donors are displayed.

### NHC antiviral activity is associated with increased viral mutation rates

It has recently been shown that NHC treatment increases the mutation rate in viral genomic RNA of RSV^24^, VEEV^11^, influenza^24^, and our previous study used RNA seq to show that overall transition mutation frequency is increased during NHC treatment of MHV and MERS-CoV during infection in continuous cell lines^13^. We sought to determine if NHC would increase the mutation frequency during MERS-CoV infection in human primary human airway epithelial cells (HAE). Using MERS-CoV infected HAE treated with either vehicle or a dose range of NHC or RDV, we show that both drugs reduced virus titers in a dose-dependent manner (Fig. 4A). We then employed a highly-sensitive high-fidelity deep sequencing approach (Primer ID NGS), which uses barcoded degenerate primers and Illumina indexed libraries to determine accurate mutation rates after antiviral treatment on viral RNA production^25^. Using this approach, we analyzed a 538bp region of viral genomic RNA in nonstructural protein 15 (nsp15). The error rates (#mutations/10,000 bases) in vehicle (0.01) or RDV (0.01) treated cultures were very low. RDV is reported to act via chain termination of nascent viral RNA, and thus the low error rates in RDV-treated cultures are in line with the proposed MOA^26^. In contrast, the error rate was significantly increased in NHC-treated MERS-CoV RNA in a dose-dependent manner (10-fold at 10μM and 5-fold at 1μM) at both 24 and 48hpi (Fig. 4C). The magnitude of the error rate in NHC-treated cultures correlated with virus titer reduction. At 48hpi the respective error rate and virus titer was 0.015 and 3.96E+06 pfu/mL for vehicle treatment, 0.045 and 2.86E+04 pfu/mL with 1μM NHC; and 0.090 and 1.5E+02 pfu/mL 10μM NHC. Thus, with 1μM NHC a 3-fold increase in error rate resulted in a 138-fold decrease in virus titer, while with 10μM NHC a 6-fold increase in error rate resulted in a 26,000-fold decrease in virus titer.

**Figure 4:**
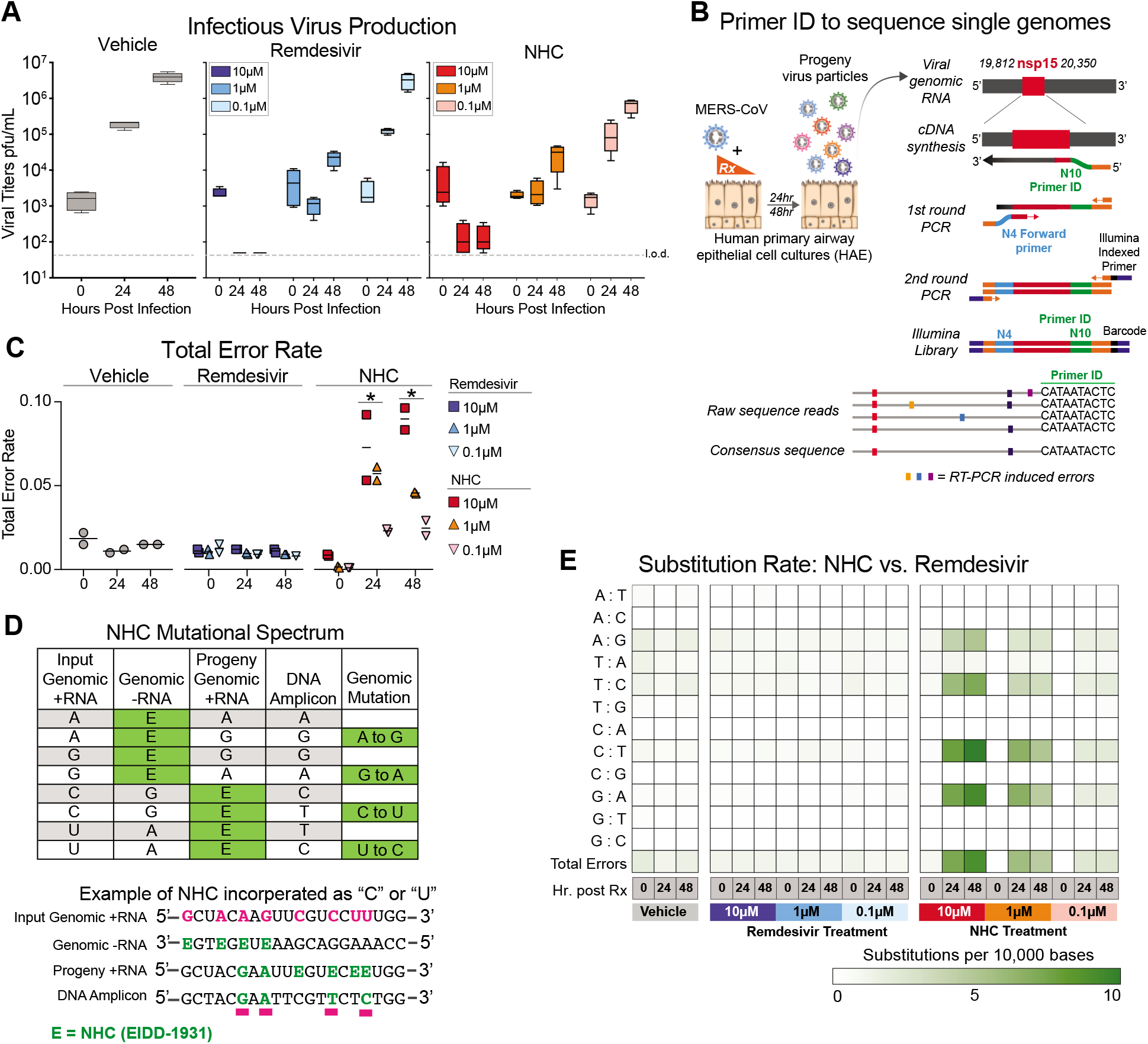
NHC antiviral activity is associated with increased viral mutation rates. **a**, Both remdesivir (RDV) and NHC reduce MERS-CoV infectious virus production in primary human HAE. Cultures were infected with MERS-CoV red fluorescent protein (RFP) at an MOI of 0.5 in duplicate in the presence of vehicle, RDV or NHC for 48 hours, after which apical washes were collected for virus titration. Data is combined from two independent studies. **b,** A deep sequencing approach called Primer ID to gain accurate sequence data for single RNA genomes of MERS-CoV. **c**, The total error rate for MERS-CoV RNA isolated from cultures in panel **a** as determined by Primer ID. Error rate values are # mutations per 10,000 bases. Asterisks indicate significant differences as compared to untreated by 2-way ANOVA with a Dunnett’s multiple comparison test. **d**, description of potential NHC mutational spectra on both positive and negative sense viral RNA. **e**, Nucleotide transitions adenine (A) to guanine (G) and uridine (U) to cytosine (C) transitions are enriched in MERS-CoV genomic RNA in an NHC dose dependent manner.

We then examined the mutational spectra induced by NHC, which can be incorporated into viral RNA as a substitution for either cytosine (C) or Uridine (U). RNA-mutagenic antivirals may incorporate in both nascent negative and positive sense RNA during genome replication (Fig. 4D). Adenine-to-guanine (A-to-G) and uracil-to-cytosine (U-to-C) transitions were enriched in MERS-CoV genomic RNA in an NHC dosedependent manner (Fig. 4E). Collectively, these data used high-fidelity sequence analysis to demonstrate a specific enrichment for A:G and C:U transitions in MERS-CoV RNA after NHC treatment of primary HAE cell cultures.

### Therapeutic EIDD-2801 reduces SARS-CoV replication and pathogenesis

Given the promising antiviral activity of NHC in vitro, we next evaluated its *in vivo* efficacy using EIDD-2801, an orally bioavailable prodrug of NHC (b-D-N^4^-hydroxycytidine-5’-isopropyl ester), designed for improved *in vivo* pharmacokinetics and oral bioavailability in humans and non-human primates^12^. Importantly, the plasma profiles of NHC and EIDD-2801 were similar in mice following oral delivery^12^. We first performed a prophylactic dose escalation study in C57BL/6 mice where we orally administered vehicle (10% PEG, 2.5% Cremophor RH40 in water) or 50, 150 or 500 mg/kg EIDD-2801 2hr prior to intranasal infection with 5E+04 PFU of mouse-adapted SARS-CoV (SARS-MA15), and then every 12hr thereafter. Beginning on 3dpi and through the end of the study, body weight loss compared to vehicle treatment was significantly diminished (50mg/kg) or prevented (150, 500mg/kg) with EIDD-2801 prophylaxis (P < 0.0001) (Supplemental Figure 2A). Lung hemorrhage was also significantly reduced 5dpi with 500mg/kg EIDD-2801 treatment (Supplemental Figure 2B). Interestingly, there was a dose-dependent reduction in SARS-CoV lung titer (median titers: 50mg/kg = 7E+03 pfu/mL, 150mg/kg = 2.5E+03 pfu/mL, 500mg/kg = 50 pfu/mL, vehicle = 6.5E+04 pfu/mL) with significant differences among the vehicle, 150 mg/kg (P = 0.03) and 500mg/kg (P = 0.006) groups. Thus, prophylactic orally administered EIDD-2801 was robustly antiviral and able to prevent SARS-CoV replication and disease.

Since only the 500mg/kg group significantly diminished weight loss, hemorrhage and reduced lung titer to near undetectable levels, we tested this dose under therapeutic treatment conditions to determine if EIDD-2801 could improve the outcomes of an ongoing CoV infection. As a control, we initiated oral vehicle or EIDD-2801 2hr prior to infection with 1E+04 pfu SARS-MA15. For therapeutic conditions, we initiated EIDD-2801 treatment 12, 24, or 48hr after infection. After initiating treatment, dosing for all groups was performed every 12hr for the duration of the study. Both prophylactic treatment initiated 2hr prior to infection and therapeutic treatment initiated 12hr after infection significantly prevented body weight loss following SARS-CoV infection on 2dpi and thereafter (−2hr: P = 0.0002 to <0.0001; +12hr: P = 0.0289 to <0.0001) (Fig. 5A). Treatment initiated 24hpi also significantly reduced body weight loss (3-5dpi, P = 0.01 to <0.0001) although not to the same degree as the earlier treatment initiation groups. When initiated 48hpi, body weight loss was only different from vehicle on 4dpi (P = 0.037, Fig. 5A). Therapeutic EIDD-2801 significantly reduced lung hemorrhage when initiated up to 24hr after infection mirroring the body weight loss phenotypes (Fig. 5B). Interestingly, all EIDD-2801 treated mice had significantly reduced viral loads in the lungs even in the +48hr group (Fig. 5C), which experienced the least protection from body weight loss and lung hemorrhage. We also measured pulmonary function via whole body plethysmography (WPB). In Figure 5D, we show the WBP PenH metric, which is a surrogate marker for bronchoconstriction or pulmonary obstruction^27^, was significantly improved throughout the course of the study if treatment was initiated up to 12hr after infection, although the +24hr group showed sporadic improvement as well (Fig. 5D). Lastly, we blindly evaluated hematoxylin and eosin stained lung tissue sections for histological features of ALI using two different and complementary scoring tools^15^, which show that treatment initiated up to +12hr significantly reduced ALI (Fig. 5E). Altogether, therapeutic EIDD-2801 was potently antiviral against SARS-CoV *in vivo* but the degree of clinical benefit was dependent on the time of initiation post infection.

**Figure 5:**
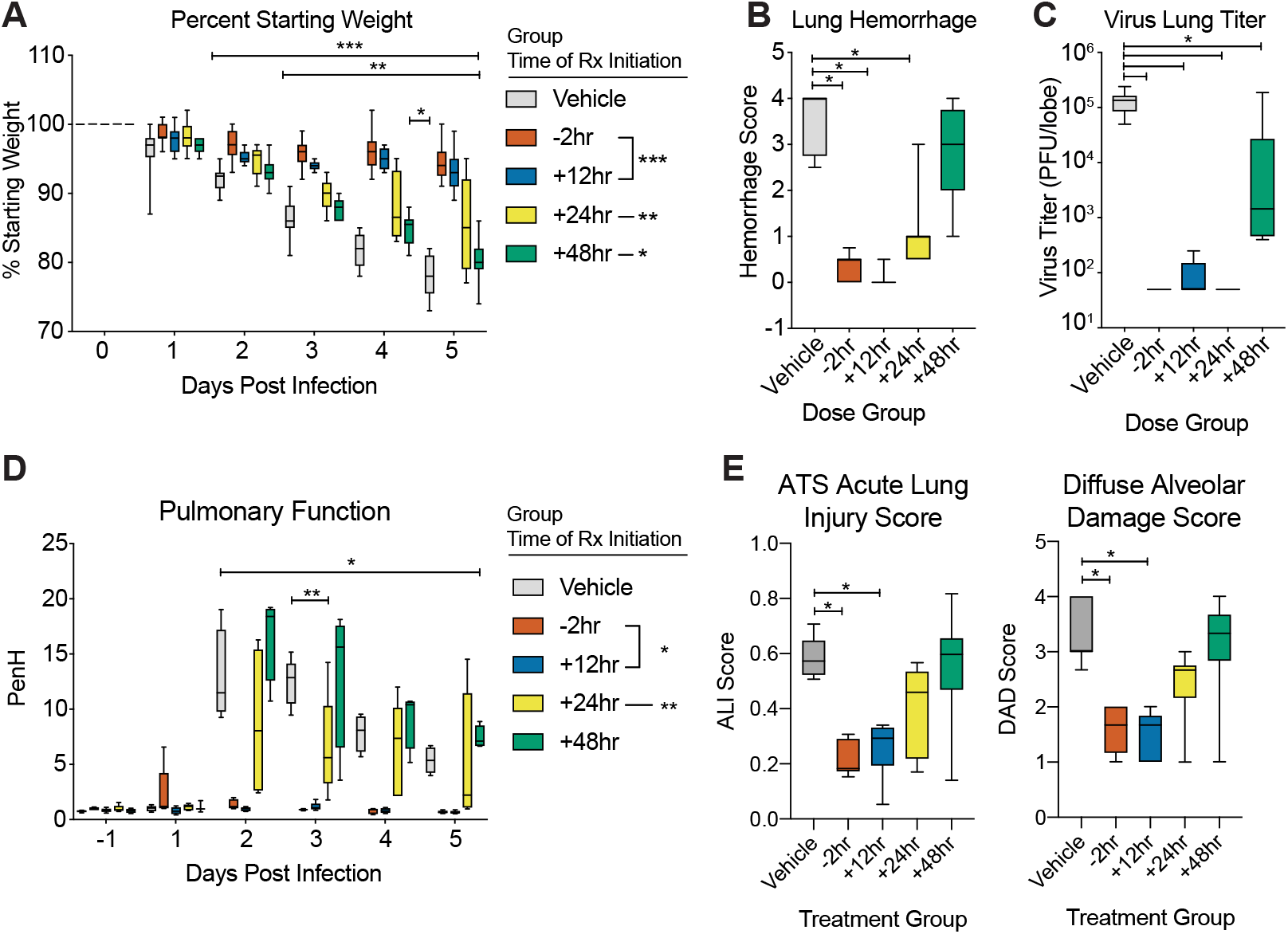
Prophylactic and Therapeutic EIDD-2801 reduces SARS-CoV replication and pathogenesis. Equivalent numbers of 25-29 week old male and female C57BL/6 mice were administered vehicle (10% PEG, 2.5% Cremophor RH40 in water) or NHC prodrug EIDD-2801 begining at −2hr, +12, +24 or +48hr post infection and every 12hr thereafter by oral gavage (n = 10/group). Mice were intranasally infected with 1E+04 PFU mouse-adapted SARS-CoV MA15 strain. **a,** Percent starting weight. Asterisks indicate differences by two-way ANOVA with Tukey’s multiple comparison test. **b,** Lung hemorrhage in mice from panel **a** scored on a scale of 0-4 where 0 is a normal pink healthy lung and 4 is a diffusely discolored dark red lung. **c**, Virus lung titer in mice from panel **a** as determined by plaque assay. Asterisks in both panel **b** and **c** indicate differences by one-way ANOVA with a Dunnett’s multiple comparison test. **d,** Pulmonary function by whole body plethysmography was performed daily on five animals per group. Asterisks indicate differences by two-way ANOVA with a Dunnett’s multiple comparison test. **e,** Therapeutic EIDD-2801 reduces acute lung injury (ALI). The histological features of ALI were blindly scored using the American Thoracic Society Lung Injury Scoring system and a Diffuse Alveolar Damage Scoring System. Three randomly chosen high power (60X) fields of diseased lung were assessed per mouse. The numbers of mice scored per group: Vehicle N = 7, −2hr N = 9, +12hr N = 9, +24hr N = 10, +48hr N = 9. Asterisks indicate statistical significance by Kruskal-Wallis with a Dunn’s multiple comparison test. For all panels, the boxes encompass the 25th to 75th percentile, the line is at the median, while the whiskers represent the range.

### Prophylactic and therapeutic EIDD-2801 reduces MERS-CoV replication and pathogenesis

After obtaining promising *in vivo* efficacy data with SARS-CoV, we investigated whether EIDD-2801 would be effective against MERS-CoV. As the murine ortholog of the MERS-CoV receptor, dipeptidyl peptidase 4 (DPP4), does not support viral binding and entry, all *in vivo* studies were performed in genetically modified mice encoding a murine DPP4 receptor encoding two human residues at positions 288 and 330 (hDPP4 288/330 mice)^15,28^. Similar to our SARS-CoV data (Supplementary Figure 3), all doses of prophylactic EIDD-2801 (50, 150 and 500mg/kg) protected hDPP4 288/330 mice from significant body weight loss (P = 0.03 to < 0.0001), lung hemorrhage (P = 0.01 to <0.0001), and virus replication which was undetectable (P < 0.0001) regardless of drug dose following intranasal infection with 5E+04 PFU mouse-adapted MERS-CoV (Supplementary Figure 4).

We then evaluated the therapeutic efficacy EIDD-2801 following the promising results of our prophylactic studies. Similar to our SARS-CoV study, EIDD-2801 treatment administered before or 12hr after intranasal mouse-adapted MERS-CoV infection (5E+04 PFU) prevented body weight loss from 2 through 6dpi (Fig. 6A, P =0.02 to <0.0001) and lung hemorrhage on 6dpi (Fig. 6B, P = 0.0004 to < 0.0001), but treatment initiated 24 or 48hr did not offer similar protection. Unlike body weight loss and lung hemorrhage data which varied by treatment initiation time, virus lung titer on 6dpi was significantly reduced to the limit of detection in all treatment groups (Fig. 6C, P < 0.0001). Interestingly, when viral genomic RNA was quantified in paired samples of lung tissue, EIDD-2801 significantly reduced levels of viral RNA (P <0.0001 to 0.017) in an initiation time-dependent manner for all groups except for +48hr (Fig. 6D). The discrepancy among infectious titers and viral RNA suggests that accumulated mutations render the particles non-infectious and undetectable by plaque assay consistent with the MOA. To gauge the effect of EIDD-2801 treatment on lung function, we assessed pulmonary function by WBP. Mirroring the body weight loss data, normal pulmonary function was only observed in groups where treatment was initiated prior to or 12hr after infection (Fig. 6E). Collectively, these data demonstrate that NHC prodrug, EIDD-2801, robustly reduces MERS-CoV infectious titers, viral RNA, and pathogenesis under both prophylactic and therapeutic conditions.

**Figure 6:**
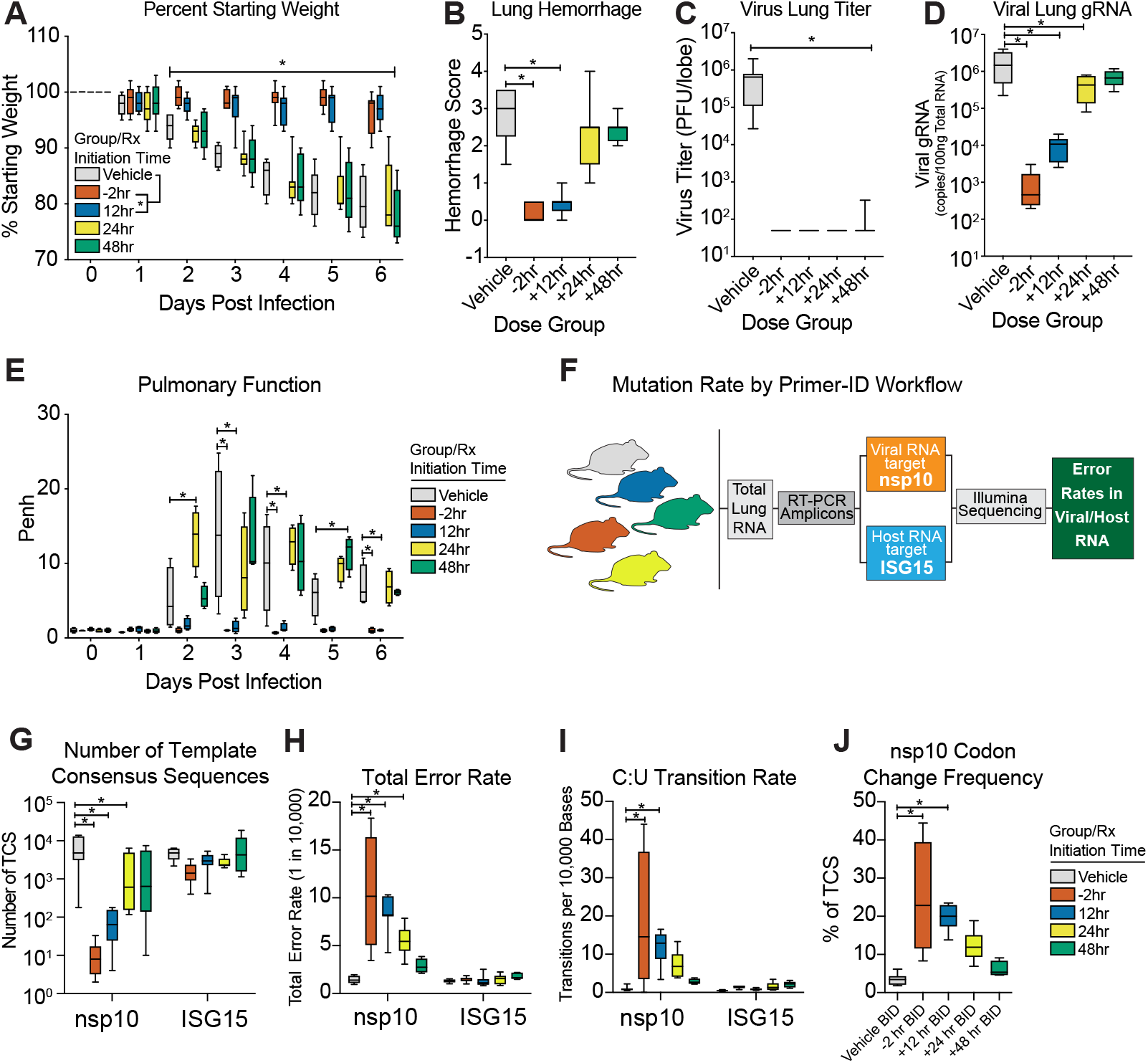
Prophylactic and therapeutic EIDD-2801 reduces MERS-CoV replication and pathogenesis coincident with increased viral mutation rates. Equivalent numbers of 10-14 week old male and female C57BL/6 hDPP4 mice were administered vehicle (10% PEG, 2.5% Cremophor RH40 in water) or NHC prodrug EIDD-2801 beginning at −2hr, +12, +24 or +48hr post infection and every 12hr thereafter by oral gavage (n = 10/group). Mice were intranasally infected with 5E+04 PFU mouse-adapted MERS-CoV M35C4 strain. **a,** Percent starting weight. Asterisks indicate differences by two-way ANOVA with Tukey’s multiple comparison test. **b,** Lung hemorrhage in mice from panel **a** scored on a scale of 0-4 where 0 is a normal pink healthy lung and 4 is a diffusely discolored dark red lung. **c**, Virus lung titer in mice from panel **a** as determined by plaque assay. Asterisks in both panel **b** and **c** indicate differences by Kruskal-Wallis with Dunn’s multiple comparison test. **d**, MERS-CoV genomic RNA in lung tissue by qRT-PCR. Asterisks indicate differences by one-way ANOVA with a Dunnett’s multiple comparison test. **e**, Pulmonary function by whole body plethysmography was performed daily on four animals per group. Asterisks indicate differences by two-way ANOVA with Tukey’s multiple comparison test. **f**, Workflow to measure mutation rate in MERS-CoV RNA and host transcript ISG15 by Primer ID in mouse lung tissue. **g**, Number of template consensus sequences for MERS-CoV nsp10 and ISG15. **h**, Total error rate in MERS-CoV nsp10 and ISG15. **i**, The cytosine to uridine transition rate in MERS-CoV nsp10 and ISG15. In panels **g-i,** asterisks indicate differences by two-way ANOVA with Tukey’s multiple comparison test. **j**, Codon change frequency in MERS-CoV nsp10. Asterisks indicate differences on Kruskal-Wallis with Dunn’s multiple comparison test.

### Therapeutic efficacy correlates with an increased MERS-CoV mutation rate *in vivo*, without increased mutations in cellular RNA

To study the molecular mechanisms associated with drug performance *in vivo*, we investigated the correlation between infectious virus production and EIDD-2801-mediated mutagenesis of MERS-CoV RNA under therapeutic treatment conditions. Using Primer ID NGS, we measured the mutation rates of both viral genomic RNA (i.e. non-structural protein 10, nsp10) and host interferon stimulated gene 15 (ISG15) mRNA, a highly upregulated innate immune related gene after MERS-CoV infection (Fig. 6F). Primer ID NGS measures the mutational frequency in single RNA molecules each of which are represented by a single template consensus sequence (TCS, See Fig. 4B)^25^. Viral TCS were significantly reduced in a treatment initiation time-dependent manner (Fig. 6G) similar to viral genomic RNA measured by qRT-PCR (Fig. 6D). In contrast, the numbers of ISG15 TCS were similar (P = 0.2 to 0.8) for all groups indicating that neither vehicle nor drug treatment significantly affected the levels of or mutated ISG15 mRNA transcripts (Fig. 6G). Similar to our TCS data in Figure 6G, the total error rate in viral nsp10 was significantly increased in groups where treatment was initiated prior to (−2hr, median error rate = 10.5 errors/10,000 bases, P < 0.0001) and up to 24hr post infection (12hr, median error rate = 8.2 errors/10,000 bases, P < 0.0001; +24hr, median error rate = 5.4 errors/10,000 bases, P = 0.0003) but the error rates in ISG15 remained at baseline for all groups (Fig. 6H). In addition, nucleotide transitions observed in MERS-CoV genomes in vitro (i.e. C to U transitions, Fig. 3), were also observed *in vivo* in groups where treatment was initiated prior to and up to 12hr post infection (P = 0.0003 to < 0.0001) (Fig. 5I). Importantly, these transitions were not observed in host ISG15 mRNA (Fig. 5I). Lastly, the EIDD-2801 dose-dependent mutagenesis of viral RNA correlated with an increase in codon change frequency, including stop codons, in mice where treatment was initiated 12hr or before (vehicle median = 3.4; −2hr median = 22.8, P = 0.0035; +12hr median = 20.0, P = 0.0004, Fig. 5I). Thus, approximately 20% of the mutations observed in the −2hr and +12hr groups resulted in a codon change and alteration of the nsp10 protein sequence. When extrapolating our results from nsp10 to the entirety of the 30kb MERS-CoV genome, EIDD-2801 likely causes between 15 (+24hr treatment) and 30 (−2hr treatment) mutations per genome 10-20% of which result in amino acid coding changes. Altogether, our data demonstrates that EIDD-2801-driven mutagenesis correlates well with the reductions in viral load, strongly suggestive of an error catastrophe-driven mechanism of action under therapeutic conditions.

## Discussion

In the past 20 years, three novel human coronaviruses have emerged^29,30^. The group 2b SARS-like CoV represent an existential and future threat to global health as evidenced by the emergence of SARS-CoV and SARS-CoV2 and zoonotic SARS-like bat CoV strains that can use human ACE2 receptors, grow well in primary human airway cells and vary by as much as 25% in key therapeutic and vaccine gene targets^8,31^. Thus, to address the current public health emergency of COVID-19 and to maximize pandemic preparedness in the future, broad-based vaccines and therapeutics, which are active against the higher risk RNA virus families prone to emergence are desperately needed.

We recently reported the broad-spectrum potency of the nucleoside prodrug, remdesivir (RDV), against an array of epidemic, contemporary and zoonotic CoV both in vitro and *in vivo*^15,16,21,31^. Currently RDV therapeutic efficacy is under investigation in several human clinical trials in China, the United States and elsewhere^32^. Here, we report the broad-spectrum antiviral activity of NHC and its orally bioavailable prodrug EIDD-2801, against SARS-CoV, MERS-CoV and related bat-CoV in primary human airway epithelial cells, as well as against the current pandemic strain SARS-CoV-2. In addition, NHC is broadly active against multiple genetically distinct viruses including coronaviruses, Venezuelan equine encephalitis (VEE), influenza A and B, Ebola, and Chikungunya viruses^10–13,16,21,24,33–35^. Here, we show that prophylactic and therapeutic EIDD-2801 significantly reduced lung viral loads and improved pulmonary function in mouse models of both SARS- and MERS-CoV pathogenesis. Although the improvement in both SARS- and MERS-CoV outcomes diminished with the increase of treatment initiation time, it is important to note that the kinetics of disease in mice are compressed as compared to that in humans. While SARS- and MERS-CoV lung titers peak on 2dpi in mice concurrent with the onset of clinical signs and notable damage to the lung epithelium, in humans this occurs 7-10 days after the onset of symptoms^16,28,36,37^. Thus, in mice, the window within which to treat emerging CoV infection prior to peak replication is compressed (e.g., 24-48hr) but should be much longer in humans. Although speculative, the SARS- and MERS-CoV *in vivo* data provided herein suggest that 2019-nCoV will prove highly vulnerable to NHC treatment modalities *in vivo*, critical experiments that must be performed as animal models become available. The data provided in this manuscript suggest that NHC should be quickly evaluated in primate models of human disease, using immediate models for MERS-CoV and SARS-CoV pathogenesis^38,39^.

Small molecule antivirals can exert their antiviral effect through multiple mechanisms including blocking viral entry, inhibiting a virally encoded enzyme, blocking virus particle formation, or targeting a host factor required for replication^40^. For VEE, EIDD-2801 exerts its antiviral activity on the RNA-dependent RNA polymerase leading to error catastrophe by inducing an error rate of replication that surpasses the error threshold allowed to sustain a virus population^11,12^. This process occurs when NHC is incorporated during RNA synthesis then subsequently misread thus increasing mutation rates. Therefore, the NHC MOA would appear less likely to be affected by the RNA proofreading activity encoded by the nsp14 exonuclease function that otherwise limits misincorperation^41^. Here, we present data using Primer ID NGS, a state of the art deep sequencing-based approach, to quantitate the frequency and identity of the mutational spectra in the MERS-CoV genome in both drug treated primary human airway cells and in mice at single genome resolution^25^. As CoV are positive sense RNA viruses that replicate through a negative sense RNA intermediate, NHC incorporation as a “C” or a “U” can occur in both polarities of RNA. Using Primer ID NGS, we found increased nucleotide transitions (A to G, G to A, C to U, U to C) consistent with those reported after influenza and VEE infections^11,12^. Under identical conditions, remdesivir did not alter the mutation rate in MERS-CoV genomic RNA, supporting its reported mechanism of action as a chain terminator of viral RNA synthesis^26^. In primary human lung cell cultures and mice infected with MERS-CoV, the NHC mutation rates inversely correlated with a reduction in infectious virus. In addition, we found a positive correlation between increased mutation rates and the frequency of nonsynonymous mutations and the degree of therapeutic efficacy in mice. To explore the potential off-target effect in host mRNA which may contribute to drug toxicity, we also examined the impact of NHC treatment on transcripts from the highly MERS-CoV induced interferon stimulated gene 15 (ISG15). While ISG15 is present in great abundance, an accumulation of mutations was not observed in ISG15 in this model even at 500mg/kg dosing. These data also support previous studies using RNAseq to demonstrate that the model coronavirus MHV displayed increased mutation frequencies following NHC treatment in vitro^13^. All together, these data strongly support the notion that EIDD-2801 and its active nucleoside analog NHC, exert their antiviral effect through the induction of error catastrophe in the targeted virus. While our data suggest that the MERS-CoV nsp14 proofreading activity appeared ineffective against NHC in vitro and EIDD-2801 *in vivo*, future studies should investigate the antiviral activity of NHC in the presence or absence of the nsp14 proofreading activity, as loss of this activity increased the sensitivity of MHV and SARS-CoV replication to remdesivir treatment^41^.

Together, our data support the continued development of EIDD-2801 as a potent broad spectrum antiviral that could be useful in treating contemporary, newly emerged and emerging coronavirus infections of the future.

## Materials and Methods

### Compounds

The parental compound b-D-N^4^-hydroxycytidine (NHC, all in vitro studies) and its prodrug EIDD-2801 (all *in vivo* studies) was supplied by Emory University Institute for Drug Discovery (EIDD). NHC was supplied as a 10mM stock in DMSO and EIDD-2801 as a solid and solubilized in vehicle containing 10% PEG400, 2.5% Cremophor RH40 in water (10/2.5/87.5%, all v/v) prior to use. Remdesivir (RDV) was solubilized in 100% DMSO and provided by Gilead Sciences, Inc as previously described^15,16^.

### Virus strains

All viruses used for these studies were derived from infectious clones and isolated as previously described^42^. Virus strains for in vitro experiments include SARS-CoV expressing the green fluorescent protein (GFP) in place of open reading frames 7a/b (ORF7a/b, SARS-GFP)^42^, bat-spike receptor binding domain (Bat-SRBD)^22^ is a chimeric CoV strain derived from the HKU3 SARS-like bat coronavirus genomic sequence that has the wild type (Urbani SARS-CoV strain) RBD in the HKU3 spike gene to allow for virus replication in non-human primate cell lines and HAE cultures, SHC014 SARS-like bat coronavirus^8^, MERS-CoV expressing nanoluciferase in the place of ORF3 (MERS-nLUC)^16^ and MERS-CoV expressing the red fluorescent protein gene in the place of ORF 5 (RFP, MERS-RFP)^43^. The virus stock utilized for MERS-CoV *in vivo* studies was derived from a plaque purified isolate of the mouse-adapted MERS-CoV p35C4 strain^44^. The virus stock utilized for SARS-CoV *in vivo* studies was derived from the infectious clone of the mouse-adapted SARS-CoV MA15 (MA15) strain^45^.

### *In vitro* experiments <u>Calu3</u>

At 48hrs prior to infection, Calu3 2B4 cells were plated in a 96-well black walled clear bottom plate at 5×10^4^ cells/well. A 10mM stock of NHC was serially diluted in 100% DMSO in 3-fold increments to obtain a ten-point dilution series. MERS-nLUC was diluted in DMEM supplemented with 10% FBS, and 1% Antibiotic-Antimycotic to achieve a multiplicity of infection (MOI) of 0.08. Cells were infected and treated with NHC in triplicate per drug dilution for 1hr, after which viral inoculum was aspirated, cultures were rinsed once and fresh medium containing drug or vehicle was added. At 48hrs post infection, nanoluciferase expression as a surrogate for virus replication was quantitated on a Spectramax (Molecular Devices) plate reader according to the manufacturer’s instructions (Promega, NanoGlo). For our 100% inhibition control, diluted MERS-nLUC was exposed to short-wave UV light (UVP, LLC) for 6 minutes to inhibit the ability of the virus to replicate. For our 0% inhibition control, cells were infected in the presence of vehicle only. DMSO was kept constant in all conditions at 0.05%. Values from triplicate wells per condition were averaged and compared to controls to generate a percent inhibition value for each drug dilution. The IC50 value was defined as the concentration at which there was a 50% decrease in luciferase expression. Data was analyzed using GraphPad Prism 8.0 (La Jolla, CA). The IC50 values were calculated by non-linear regression analysis using the doseresponse (variable slope) equation (four parameter logistic equation): Y = Bottom + (Top-Bottom)/(1+10^((LogIC50-X)*HillSlope)). To measure cell viability to determine if there was any NHC induced cytotoxicity, Calu3 2B4 cells were plated and treated with NHC only as described above. Cells were exposed to the same ten-point dilution series created for the in vitro efficacy studies. As above, 0.05% DMSO-treated cells served as our 0% cytotoxicity control. Wells without cells served as our 100% cytotoxic positive control. After 48hr, cell viability was measured on a Spectramax (Molecular Devices) via Cell-Titer Glo Assay (Promega) according to the manufacturer’s protocol. Similar data was obtained in three independent experiments.

### HAE

Human tracheobronchial epithelial cells were obtained from airway specimens resected from patients undergoing surgery under University of North Carolina Institutional Review Board-approved protocols by the Cystic Fibrosis Center Tissue Culture Core. Primary cells were expanded to generate passage 1 cells and passage 2 cells were plated at a density of 250,000 cells per well on Transwell-COL (12mm diameter) supports. Human airway epithelium cultures (HAE) were generated by provision of an air-liquid interface for 6 to 8 weeks to form well-differentiated, polarized cultures that resembled *in vivo* pseudostratified mucociliary epithelium^46^. At 48 hours prior to infection the apical surface of the culture was washed with 500 μL PBS for 1.5 hours at 37°C and the cultures moved into fresh air liquid interface (ALI) media. Immediately prior to infection, apical surfaces were washed twice with 500 μL of PBS with each wash lasting 30 minutes at 37°C and HAE cultures were moved into ALI media containing various concentrations of NHC ranging from 10 μM to 0.0016 μM as indicated for each experiment (final % DMSO < 0.05%). Upon removing the second PBS wash, 200 μL of viral inoculum (multiplicity of infection of (MOI) 0.5) was added to the apical surface and HAE cultures were incubated for 3 hours at 37°C. Viral inoculum was then removed, and the apical surface of the cultures were washed three times with 500μL PBS and then incubated at 37°C until 48 hours post infection (hpi). For all HAE cultures, infectious virus produced was collected by washing the apical surface of the culture with 100 μL PBS. Apical washes were stored at −80 °C until analysis and titered by plaque assay as previously described^16^.

### qRT-PCR approach to assess cytotoxicity

Total RNA was isolated using the Zymo Direct-zol RNA MiniPrep Kit (Zymo Research Corp., Irvine, CA, USA) according to the manufacturer’s directions. First-strand cDNA was generated using Superscript III reverse transcriptase (Life Technologies, Carlsbad, CA, USA). For quantification of cellular markers of toxicity/apoptosis, real-time PCR was performed using commercially validated TaqMan-based primer-probe sets (**Supplementary Table 1**) and TaqMan Universal PCR Mix (Life Technologies). Results were then normalized as described above.

### Primer ID and Deep Sequencing

Primer ID NGS is designed to specifically identify and remove RT-PCR mutations, while facilitating highly accurate sequence determination of single RNA molecules, because each cDNA is created with a barcoded degenerate primer (N10, 4^10^ combinations) from which Illumina indexed libraries are made. We used a multiplexed Primer ID library prep approach and MiSeq sequencing to investigate the presence of mutations in the viral genomes and murine mRNA. We designed cDNA primers targeting multiple regions on the viral genome and murine mRNA, each with a block of random nucleotides (11 bp long) as the Primer ID^25,47^ (**Supplementary Table 2**). Viral RNA was extracted using QIAamp viral RNA kit. A pre-amplification titration of templates was performed to estimate the amount of template to use. We used SuperScript III to make cDNA with multiplexed cDNA primers based on the regions to be sequenced. We used 41R_PID11 for the pilot sequencing and titration determination. For the MERS-CoV sequencing, we multiplexed nsp10_PID11, nsp12_PID11 and nsp14_PID11 for the cDNA reaction; for the murine mRNA sequencing, we used mixed primers of nsp10_PID11, ifit3_PID11, isg15_PID11. After bead purification, we amplified the cDNA with a mixture of forward primers (based on the described schemes) and a universal reverse primer, followed by another round of PCR to incorporate Illumina sequencing adaptors and barcodes in the amplicons. After gel-purification and quantification, we pooled 24 libraries for MiSeq 300 base paired-end sequencing. The TCS pipeline version 1.38 (https://github.com/SwanstromLab/PID) was used to process the Primer ID sequencing data and construct template consensus sequences (TCSs) to represent each individual input templates, and the sequences of each region in the pool was de-multiplexed. The RUBY package viral_seq version 1.0.6 (https://rubygems.org/gems/viral_seq) was used to calculate the mutation rate at each position.

### *In vivo* experiments

We performed 4 mouse studies to evaluate the *in vivo* efficacy of the NHC prodrug (EIDD-2801). First, we performed prophylactic dose escalation studies for both SARS- and MERS-CoV to determine the most efficacious dose of EIDD-2801 per virus. For SARS-CoV, in cohorts of equivalent numbers of male and female 20-29 week old SPF C57BL/6J (Stock 000664 Jackson Labs) mice (n = 10/dose group), we administered vehicle (10% PEG, 2.5% Cremophor RH40 in water) or 50, 150 or 500mg/kg EIDD-2801 by oral gavage 2hr prior to intranasal infection with 1E+04 PFU mouse-adapted SARS-CoV strain MA15 in 50μl. Mice were anaesthetized with a mixture of ketamine/xylazine prior to intranasal infection. Vehicle or drug was administered every 12hr for the remainder of the study. Body weight and pulmonary function by whole body plethysmography was measured daily. On 5dpi, animals were sacrificed by isoflurane overdose, lungs were scored for lung hemorrhage, and the inferior right lobe was frozen at −80°C for viral titration via plaque assay. Briefly, 500,000 Vero E6 cells/well were seeded in 6-well plates. The following day, medium was removed and serial dilutions of clarified lung homogenate were added per plate (10^-1^ to 10^-6^ dilutions) and incubated at 37°C for 1hr after which wells were overlayed with 1X DMEM, 5% Fetal Clone 2 serum, 1X antibiotic/antimycotic, 0.8% agarose. Two days after, plaques were enumerated to generate a plaque/ml value. Lung hemorrhage is a gross pathological phenotype readily observed by the naked eye driven by the degree of virus replication where the coloration of the lung changes from pink to dark red^48,49^. The large left lobe was placed in 10 % buffered formalin and stored at 4°C for 1-3 weeks until histological sectioning and analysis. For MERS-CoV, the prophylactic dose escalation studies we performed exactly as done for SARS-CoV with the following exceptions. First, MERS-CoV binds the human receptor dipeptidyl peptidase 4 (DPP4) to gain entry into cells and two residues (288 and 330) in the binding interface of mouse DPP4 prevent infection of mice. We recently developed a mouse model for MERS-CoV through the mutation of mouse DPP4 at 288 and 330 thus humanizing the receptor (*hDPP4*) and rendering mice susceptible to MERS-CoV infection^28^. We performed all *in vivo* studies with EIDD-2801 in equivalent numbers of 10-14 week old female and male C57BL/6J hDPP4 mice. Second, we intranasally infected mice with 5E+04 PFU mouse-adapted MERS-CoV strain M35C4 in 50μl. Third, to titer lungs by plaque assay, Vero CCL81 cells were used and plaques were enumerated 3 days post infection.

To determine the time at which therapeutic administration of EIDD-2801 would fail to improve outcomes with SARS-CoV or MERS-CoV infection, we performed therapeutic efficacy studies in mice where we initiated treatment 2hr prior to infection or 12, 24 or 48hr after infection. As 500mg/kg provided the most complete protection from disease in prophylactic SARS-CoV studies, this dose was used for both therapeutic efficacy studies. Vehicle or EIDD-2801 was given via oral gavage twice daily following initiation of treatment. For both SARS-CoV and MERS-CoV, the infectious dose for the therapeutic studies and the mouse strains were the same as that used in the prophylactic studies. The numbers of mice per group for the SARS-CoV studies were as follows: Vehicle (n = 10), −2hr (n = 10), +12hr (n = 10), +24hr (n = 10), +48hr (n = 10). The numbers of mice per group for the MERS-CoV therapeutic studies were as follows: Vehicle (n = 9), −2hr (n = 9), +12hr (n = 9), +24hr (n = 7), +48hr (n = 10). As described above, each day mouse body weight and pulmonary function was quantitated. On 5dpi for SARS-CoV and 6dpi for MERS-CoV, animals were humanely sacrificed and tissues were harvested and analyzed as described above. In addition, for the MERS-CoV study, lung tissue was harvested and stored in RNAlater (Thermo Fisher) at −80°C and then thawed, homogenized in Trizol reagent (Invitrogen) and total RNA was isolated using a Direct-zol RNA MiniPrep kit (Zymo Research). This total RNA was then used for Primer ID and qRT-PCR.

### Whole body plethysmography

Pulmonary function was monitored once daily via whole-body plethysmography (Buxco Respiratory Solutions, DSI Inc.). Mice destined for this analysis were chosen prior to infection. Briefly, after a 30-minute acclimation time in the plethysmograph, data for 11 parameters was recorded every 2 seconds for 5 minutes.

### Acute lung injury histological assessment tools

Two different and complementary quantitative histologic tools were used to determine if antiviral treatments diminished the histopathologic features associated with lung injury. Both analyses and scoring were performed by a Board Certified Veterinary Pathologist who was blinded to the treatment groups.

### American Thoracic Society Lung Injury Scoring Tool

In order to help quantitate histological features of ALI observed in mouse models and increase their translation to the human condition, we used the ATS scoring tool^49^. In a blinded manner, we chose three random diseased fields of lung tissue at high power (60 ×), which were scored for the following: (A) neutrophils in the alveolar space (none = 0, 1–5 cells = 1, > 5 cells = 2), (B) neutrophils in the interstitial space/ septae (none = 0, 1–5 cells = 1, > 5 cells = 2), (C) hyaline membranes (none = 0, one membrane = 1, > 1 membrane = 2), (D) Proteinaceous debris in air spaces (none = 0, one instance = 1, > 1 instance = 2), (E) alveolar septal thickening (< 2× mock thickness = 0, 2–4× mock thickness = 1, > 4× mock thickness = 2). To obtain a lung injury score per field, the scores for A–E were then put into the following formula, which contains multipliers that assign varying levels of importance for each phenotype of the disease state.: score = [(20× A) + (14 × B) + (7 × C) + (7 × D) + (2 × E)]/100. The scores for the three fields per mouse were averaged to obtain a final score ranging from 0 to and including 1.

### Diffuse Alveolar Damage (DAD) Tool

The second histological tool to quantitate lung injury was reported by Schmidt et al.^50^. DAD is the pathological hallmark of ALI^49,50^. Three random diseased fields of lung tissue were score at high power (60 ×) for the following in a blinded manner: 1 = absence of cellular sloughing and necrosis, 2 = Uncommon solitary cell sloughing and necrosis (1–2 foci/field), 3 = multifocal (3 + foci) cellular sloughing and necrosis with uncommon septal wall hyalinization, or 4 = multifocal (>75% of field) cellular sloughing and necrosis with common and/or prominent hyaline membranes. The scores for the three fields per mouse were averaged to get a final DAD score per mouse.

### MERS-CoV genomic RNA qRT-PCR

Mouse lungs were stored in RNAlater (ThermoFisher) at −80°C until processed via homogenization in TRIzol (Invitrogen). Total RNA was isolated using Direct-zol RNA MiniPrep kit (Zymo Research). Previously published TaqMan primers were synthesized by Integrated DNA Technologies (IDT) to quantify MERS genomic RNA (targeting orf1a. Forward: 5’-GCACATCTGTGGTTCTCCTCTCT-3’, Probe (6-FAM/ZEN/IBFQ): 5’-TGCTCCAACAGTTACAC-3’, Reverse: 5’-AAGCCCAGGCCCTACTATTAGC)^51^. qRT-PCR was performed using 100ng total RNA compared to an RNA standard curve using TaqMan Fast Virus 1-Step Master Mix (ThermoFisher) on a Quant Studio 3 (Applied Biosystems).

### nsp12 phylogenetic analysis and conservation modeling

Coronavirus RdRp (nsp12) protein sequence alignments and phylogenetic trees were generated using Geneious Tree Builder in Geneious Prime (version 2020.0.5) and visualized using Evolview (https://www.evolgenius.info/evolview/). Protein similarity scores were calculated using Blosom62 matrix. The accession numbers used were: PDCoV (KR265858), AIBV (NC_001451), HCoV-229E (JX503060), PEDV (NC_003436), MHV (AY700211), HCoV-HKU1 (DQ415904), HCoV-NL63 (JX504050), HCOV-OC43 (AY903460), HKU5-1 (NC_009020), MERS-CoV (JX869059), HKU9-4 (EF065516), 2019-nCoV (MN996528), HKU3-1 (DQ022305), SHC014 (KC881005), WIV1 (KF367457), SARS-CoV (AY278741). Amino acid conservation scores of coronavirus RdRp were generated using ConSurf Server (https://consurf.tau.ac.il/) using the protein alignment described above and visualized on the SARS-CoV RdRp structure (PDB: 6NUR) in PyMol (version 1.8.6.0)^20,52^.

### Statistical analysis

All statistical data analyses were performed in Graphpad Prism 8. Statistical significance for each endpoint was determined with specific statistical tests. For each test, a p-value <0.05 was considered significant. Specific tests are noted in each figure legend.

### Ethics regulation of laboratory animals

Efficacy studies were performed in animal biosafety level 3 facilities at UNC Chapel Hill. All work was conducted under protocols approved by the Institutional Animal Care and Use Committee at UNC Chapel Hill (IACUC protocol #16-284) according to guidelines set by the Association for the Assessment and Accreditation of Laboratory Animal Care and the U.S. Department of Agriculture.

## Funding

We would like to acknowledge the following funding sources, Antiviral Drug Discovery and Development Center (5U19AI109680), a partnership grant from the National Institutes of Health (5R01AI132178) and an NIAID R01 grant (AI108197). NIAID contract, HHSN272201500008C, was awarded to G.P. and The Emory institute for Drug Development and a subcontract from this was awarded to R.S.B. and M.R.D.

## Author contributions

A.C.S., T.P.S, designed *in vitro* efficacy studies. A.C.S., A.J.B., T.P.S. and R.L.G. executed and/or analyzed *in vitro* efficacy studies. J.H., A.T. and N.J.T. providing the clinical isolate of SARS-CoV-2. T.P.S., A.A.K., M.G.N., G.P. and R.S.B. designed *in vivo* efficacy studies. T.P.S., A.C.S., S.Z., C.S.H, and R.S. designed, executed and/or analyzed the Primer ID NGS data. K.H.D 3^rd^ performed structural modeling and phylogenetics and sequence alignments. M.L.A., A.J.P., J.D.C. and M.R.D. designed, performed and/or executed the construction of RDV resistant MHV and performed cross-resistance studies. T.P.S., A.S. and S.R.L. executed and analyzed *in vivo* efficacy studies. A.S. and S.R.L. performed whole body plethysmography for *in vivo* studies. S.A.M. assessed all lung pathology. G.R.B., and M.S., were responsible for synthesis, and scale-up of small molecules. T.P.S., A.C.S., S.Z., S.R.L, A.S., K.H.D. 3^rd^, M.L.A., A.J.P., J.D.C, G.R.B., A.A.K., G.P., R.S. M.R.D., and R.S.B., wrote the manuscript.

## Disclaimers

The findings and conclusions in this report are those of the author(s) and do not necessarily represent the official position of the Centers for Disease Control and Prevention. Names of specific vendors, manufacturers, or products are included for public health and informational purposes; inclusion does not imply endorsement of the vendors, manufacturers, or products by the Centers for Disease Control and Prevention or the US Department of Health and Human Services.

## Competing financial interests

A.C.S. received a contract from NIAID to support the *in vitro* and *in vivo* efficacy studies reported herein. UNC is pursuing IP protection for Primer ID and R.S. has received nominal royalties.

---

**Supplementary Figure 1:**
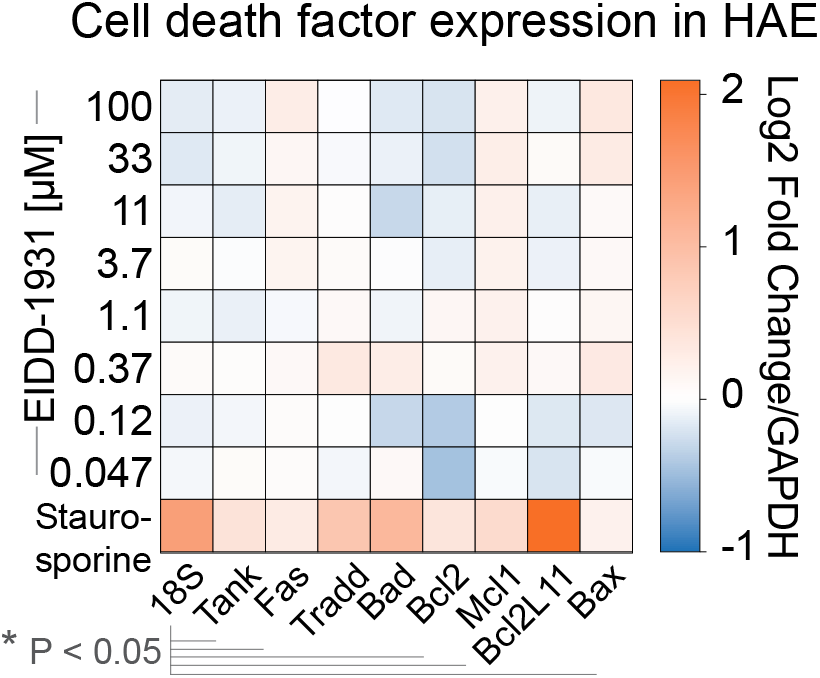
Assessment of cytotoxicity of NHC in primary human epithelial cell cultures by qRT-PCR. Companion figure to **Figure 1c** and **d**. Primary human epithelial cell cultures were exposed to positive control 1μM staurosporine or a dose response of NHC for 48hr. Cytotoxicity was assessed by qRT-PCR for cell death factor gene expression.

**Supplementary Figure 2:**
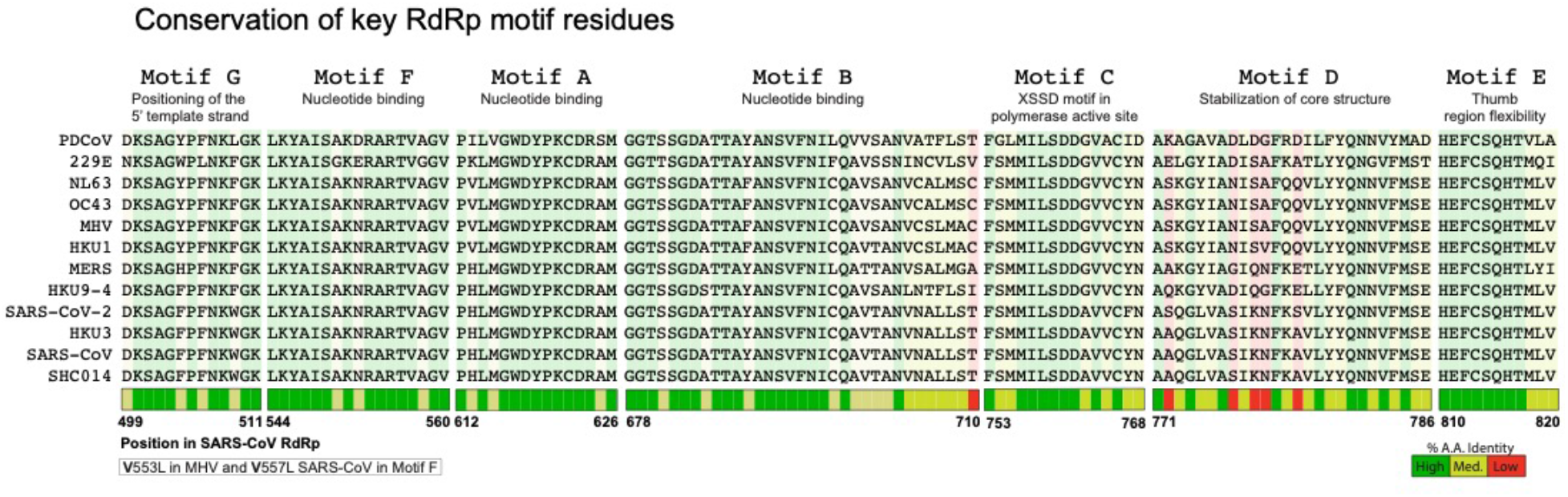
High conservation of RdRp functional domains for SARS-CoV-2. Companion figure to **Figure 2a.** Multiple sequence alignment of the RNA dependent RNA polymerase (RdRp) from viruses in the dendrogram in Figure 2a showing high conservation in the RdRp structural motifs A-G.

**Supplementary Figure 3:**
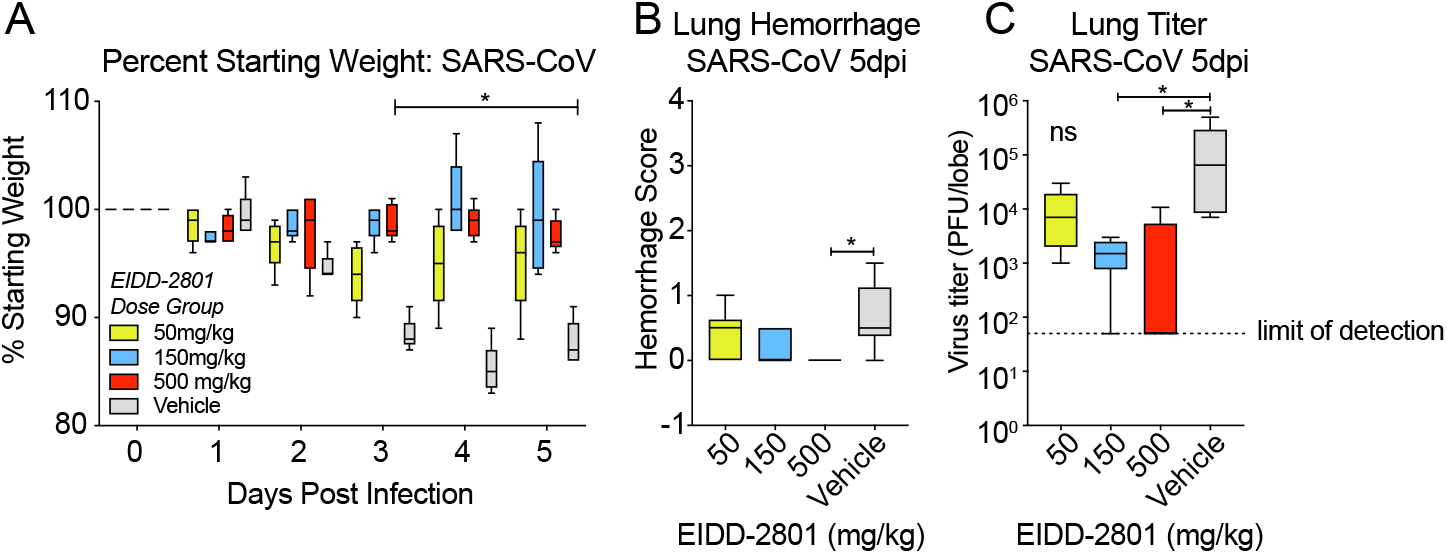
Prophylactic EIDD-2801 reduces SARS-CoV replication and pathogenesis. Companion figure to **Figure 5**. Equivalent numbers of 20 week old male and female C57BL/6 mice were administered vehicle (10% PEG, 2.5% Cremophor RH40 in water) or NHC prodrug EIDD-2801 beginning at 2hr prior to infection and every 12hr thereafter by oral gavage (n = 10/group). Mice were intranasally infected with 1E+04 PFU mouse-adapted SARS-CoV MA15 strain. **a,** Percent starting weight. Asterisks indicate differences by two-way ANOVA with Dunnett’s multiple comparison test. **b,** Lung hemorrhage in mice from panel **a** scored on a scale of 0-4 where 0 is a normal pink healthy lung and 4 is a diffusely discolored dark red lung. **c**, Virus lung titer in mice from panel **a** as determined by plaque assay. Asterisks in both panel **b** and **c** indicate differences by Kruskal-Wallis with a Dunn’s multiple comparison test.

**Supplementary Figure 4:**
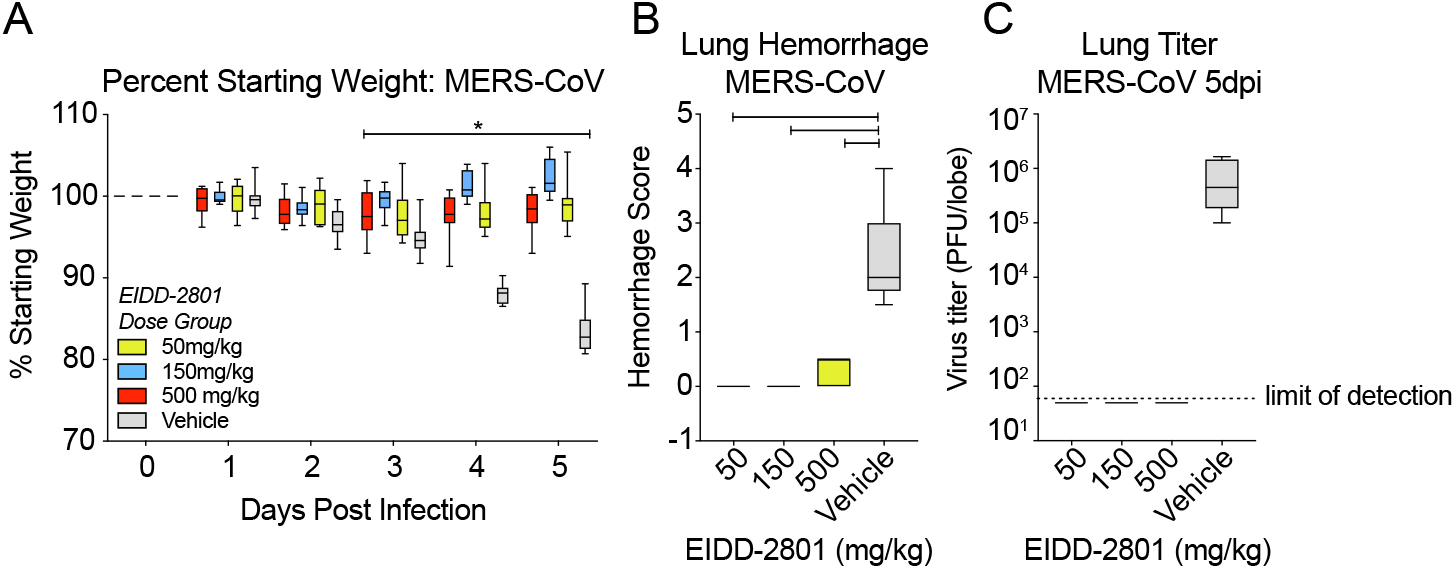
Prophylactic EIDD-2801 reduces MERS-CoV replication and pathogenesis. Companion figure to **Figure 6.** Equivalent numbers of 10-14 week old male and female C57BL/6 hDPP4 mice were administered vehicle (10% PEG, 2.5% Cremophor RH40 in water) or NHC prodrug EIDD-2801 beginning 2hr prior to infection every 12hr thereafter by oral gavage (n = 10/group). Mice were intranasally infected with 5E+04 PFU mouse-adapted MERS-CoV M35C4 strain. **a,** Percent starting weight. Asterisks indicate differences by two-way ANOVA with Dunnett’s multiple comparison test. **b,** Lung hemorrhage in mice from panel **a** scored on a scale of 0-4 where 0 is a normal pink healthy lung and 4 is a diffusely discolored dark red lung. **c**, Virus lung titer in mice from panel **a** as determined by plaque assay. Asterisks in both panel **b** and **c** indicate differences by Kruskal-Wallis with Dunn’s multiple comparison test.

**Supplementary Table 1.** Real-time PCR primer/probe sets for indicators of cellular apoptosis/toxicity

| Primer/Probe Target | Assay Reference Number* |
| --- | --- |
| Bax | Hs00180269_m1 |
| Bad | Hs00188930_m1 |
| Bcl2L11 | Hs00708019_s1 |
| Bcl2 | Hs00608023_m1 |
| Mcl1 | Hs01050896_m1 |
| Tradd | Hs00601065_g1 |
| Fas | Hs00236330_m1 |
| Tank | Hs00370305_m1 |
| 18S | 4352930E |
| GAPD** | 4352934E |

\* Validated assays available from Life Technologies
\*\* The housekeeping gene hGAPDH was used for normalization of real-time results.

**Supplementary Table 2.**
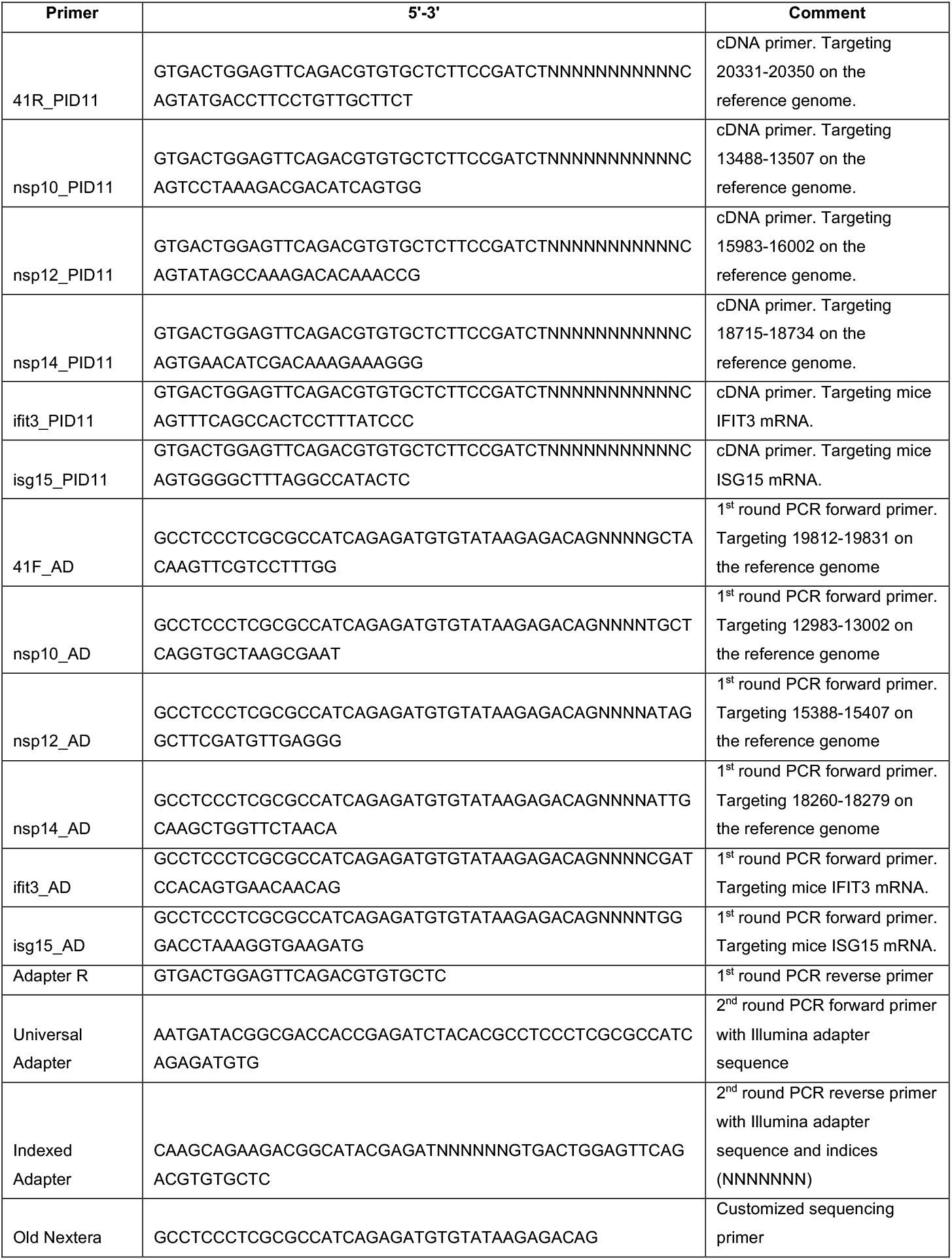
Primer used for MiSeq library prep and sequencing.

